# Male RORγt-transgenic mice exhibit lower body weight and an altered locomotor response to a novel environment

**DOI:** 10.64898/2026.08.31.748441

**Authors:** Tetsuya Sasaki, Aki Takahashi, Yosuke Takei

**Affiliations:** Laboratory of Anatomy and Neuroscience, Department of Biomedical Sciences, Institute of Medicine, University of Tsukuba, Tsukuba, Ibaraki, Japan; Laboratory of Behavioral Neurobiology, Institute of Human Sciences, University of Tsukuba, Tsukuba, Ibaraki, Japan; Master’s and Doctoral Programs in Neuroscience, Degree Programs in Comprehensive Human Sciences, Graduate School of Comprehensive Human Sciences, University of Tsukuba, Tsukuba, Ibaraki, Japan

**Author notes:** Correspondence: Tetsuya Sasaki and Yosuke Takei.

**Keywords:** RORγt, Rorc, transgenic mice, novelty response, light–dark box, resident–intruder test, locomotor activity, body weight

## Abstract

RORγt, encoded by *Rorc*, is a lineage-defining regulator of T helper 17 (Th17) cell differentiation, and IL-17A signaling has been implicated in neurodevelopment and behavior. We characterized baseline behavioral phenotypes in male RORγt-transgenic mice with T-cell-biased RORγt overexpression and wild-type (WT) littermates. Mice were assessed using the resident–intruder paradigm and the light–dark box test, followed by body-weight measurement. RORγt-transgenic mice had significantly lower body weight than WT mice. No genotype differences were detected in attack latency, attack bites, attack incidence, time in the light compartment, light–dark transitions, or transition latency. In contrast, locomotor activity analyzed in 1-min bins showed a significant genotype × time interaction.

Transgenic mice traveled nominally less during the first minute of the light–dark box test, although this single-bin comparison did not remain significant after correction for multiple comparisons. These findings indicate that T-cell-specific RORγt overexpression is associated with altered temporal dynamics of locomotor activity during initial exposure to a novel environment rather than with a generalized anxiety-like or aggression-related phenotype.

## Introduction

Interleukin-17A (IL-17A) is best known as a proinflammatory cytokine produced by Th17 and other immune-cell populations, but accumulating evidence indicates that IL-17A signaling can also influence the nervous system and behavior. Maternal IL-17A signaling can alter fetal cortical development and promote autism-like behavioral phenotypes in mouse offspring (Choi et al., 2016), whereas IL-17A acting on cortical neurons can enhance sociability in several mouse models of neurodevelopmental disorders (Reed et al., 2020). In adult mice, IL-17A derived from meningeal γδ T cells contributes to the regulation of anxiety-like behavior through neuronal IL-17 receptor signaling (Alves de Lima et al., 2020). Direct exposure of the fetal brain to IL-17A also activates cortical microglia and alters their localization (Sasaki et al., 2020). Recent reviews therefore place the RORγt–IL-17A axis within a broader framework of immune–brain communication across development and adulthood (Kubo et al., 2025; Sanaka et al., 2026).

Differentiation of Th17 cells is directed by the orphan nuclear receptor RORγt, encoded by *Rorc* (Ivanov et al., 2006). The CD2-RORγt transgenic line used here expresses full-length murine RORγt under human CD2 regulatory elements, producing preferential RORγt overexpression in T cells. This line exhibits a Th17-biased immune phenotype and elevated circulating IL-17A (Yoh et al., 2012). Previous studies using the same line have reported altered inflammatory responses during pregnancy, including enhanced susceptibility to poly(I:C)-associated fetal loss (Tome et al., 2019), as well as subtle central nervous system phenotypes (Sasaki et al., 2021). In addition, adoptive transfer of Th17 cells can promote depression-like behavior in mice (Beurel et al., 2013), further supporting the possibility that sustained changes in the RORγt/Th17 axis may influence behavioral state.

Despite these observations, the baseline behavioral phenotype associated with constitutive T-cell-specific RORγt overexpression has not been systematically characterized in the absence of an acute immune challenge. We therefore examined male RORγt-transgenic (RORγt-Tg) mice and WT littermates in two behavioral paradigms relevant to social aggression and anxiety-related behavior: the resident–intruder test and the light–dark box test. In addition to conventional light–dark indices, we analyzed the temporal profile of locomotor activity during the 10-min session to determine whether the genotypes differed in their initial response to a novel environment or in within-session activity dynamics. Body weight was measured immediately after behavioral testing.

## Results

### RORγt-Tg mice have lower body weight

Male RORγt-Tg mice and WT littermates were tested in three independent runs at 81–100 days of age and pooled for analysis (WT *n* = 12; Tg *n* = 13; Figure 1A). RORγt-Tg mice had lower body weight than WT littermates (25.7 ± 0.4 g versus 28.1 ± 0.5 g; *t*(23) = 3.69, *p* = 0.0012; Figure 1B). The genotype effect remained significant in a sensitivity model that adjusted for test run and age at testing (*p* = 0.0018). The direction of this difference is consistent with the reported role of IL-17 signaling in limiting adipogenesis and weight gain (Zúñiga et al., 2010), although IL-17A was not measured in the animals tested here.

**Figure 1.**
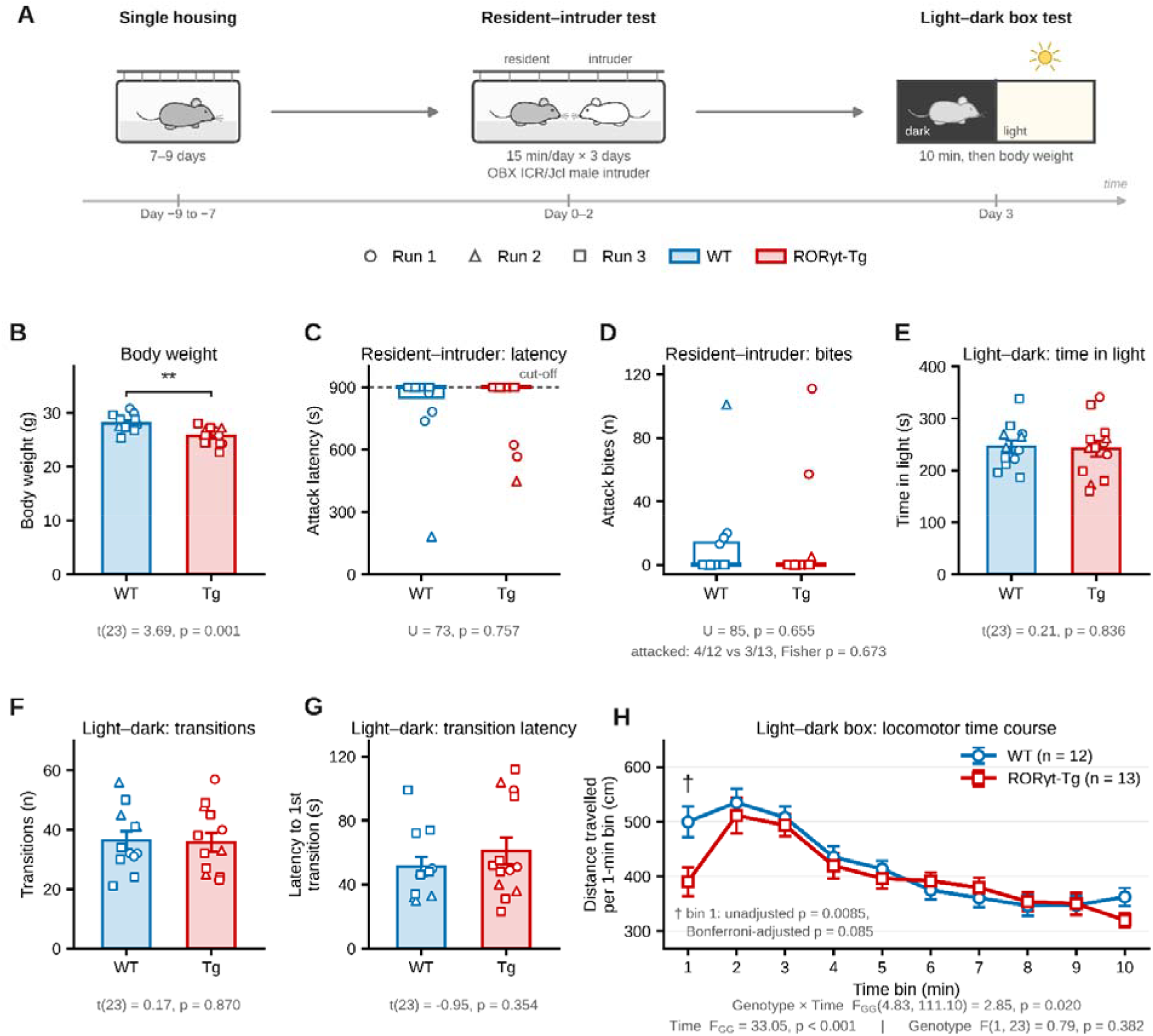
Male RORγt-transgenic mice exhibit lower body weight and an altered locomotor response to a novel environment. **(A)**Experimental design. Male RORγt-Tg mice (Tg, n = 13) and WT littermates (n = 12) were single-housed for 7–9 days, tested in the resident– intruder paradigm for 15 min/day on three consecutive days, and tested in the light–dark box for 10 min on the following day; body weight was then measured. Data were collected in three runs: run 1, 3 WT/3 Tg at 100 days; run 2, 3 WT/3 Tg at 81 days; and run 3, 6 WT/7 Tg at 93–99 days. Circles, triangles, and squares denote runs 1, 2, and 3, respectively. (B)Body weight (mean ± s.e.m.) was lower in Tg mice (t(23) = 3.69, p = 0.0012). (C, D) Resident–intruder outcomes are shown as median and interquartile range because latency was censored at 900 s and bite counts were zero-inflated. (C) Mean latency to first attack across three sessions. (D) Total attack bites. Neither outcome differed by genotype (Mann–Whitney U tests: latency U = 73, p = 0.757; bites U = 85, p = 0.655). Attack incidence was 4/12 WT and 3/13 Tg (Fisher’s exact p = 0.673); no mouse attacked in run 3. (E–G) Conventional light–dark box outcomes (mean ± s.e.m.): (E) time in light, (F) transitions, and (G) latency to first light-compartment entry. No genotype difference was detected (t(23) = 0.21, p = 0.836; t(23) = 0.17, p = 0.870; and t(23) = ™0.95, p = 0.354, respectively). (H) Distance traveled per 1-min bin (mean ± s.e.m.). Mixed-model ANOVA with Greenhouse–Geisser correction showed a genotype × time interaction (FGG(4.83, 111.10) = 2.85, p = 0.020) and a time effect (FGG(4.83, 111.10) = 33.05, p < 0.001), but no genotype effect (F(1, 23) = 0.79, p = 0.382). At minute 1, Tg mice traveled nominally less than WT mice (t(23) = 2.88, unadjusted p = 0.0085; Bonferroni-adjusted p = 0.085); other bins did not differ. Double asterisks indicate p < 0.01; † indicates nominal significance before correction. All tests were two-sided.

### Resident–intruder aggression does not differ detectably between genotypes

Aggression was assessed across three consecutive 15-min resident–intruder sessions. No genotype difference was detected in latency to the first attack (Mann–Whitney *U* = 73, *p* = 0.757) or in the total number of attack bites (*U* = 85, *p* = 0.655; Figure 1C,D). The proportion of residents that attacked at least once was also similar (4/12 WT versus 3/13 Tg; Fisher’s exact test, *p* = 0.673). No mouse of either genotype attacked in run 3, indicating that the aggression comparison was driven primarily by the first two runs. Thus, under the present conditions, the data do not provide evidence for a large genotype effect on offensive aggression, while the low baseline attack rate limits sensitivity to smaller effects.

### Conventional light–dark box measures are unchanged

The light–dark box was used to assess conventional indices of approach–avoidance behavior. Time spent in the light compartment (*t*(23) = 0.21, *p* = 0.836), the number of light–dark transitions (*t*(23) = 0.17, *p* = 0.870), and latency to the first transition into the light compartment (*t*(23) = ™0.95, *p* = 0.354) did not differ detectably between genotypes (Figure 1E–G). These measures therefore do not support a generalized anxiety-like phenotype in RORγt-Tg mice.

### RORγt-Tg mice show an altered locomotor time course during initial exposure to the apparatus

When locomotor activity was resolved into 1-min bins, both groups showed a marked decline in activity over the course of the session, consistent with within-session adaptation to the apparatus (Figure 1H). A mixed-model ANOVA showed a significant genotype × time interaction after Greenhouse–Geisser correction (*F*GG(4.83, 111.10) = 2.85, *p* = 0.020) and a robust main effect of time (*F*GG(4.83, 111.10) = 33.05, *p* < 0.001), but no main effect of genotype (*F*(1, 23) = 0.79, *p* = 0.382). Post hoc bin-wise comparisons localized the largest difference to the first minute, during which Tg mice traveled less than WT mice (*t*(23) = 2.88, unadjusted *p* = 0.0085). This contrast did not survive Bonferroni correction across the ten bins (adjusted *p* = 0.085), and no later bin differed at the unadjusted 0.05 level.

The minute-1 estimate was materially unchanged after adjustment for test run and age (*p* = 0.0095). Because RORγt-Tg mice were lighter than WT mice, we also tested whether the early locomotor difference was secondary to body weight. In a model including body weight as a covariate, the genotype term remained significant (*p* = 0.014), whereas body weight itself was not independently associated with minute-1 distance (*p* = 0.51). These analyses indicate that the overall genotype × time interaction reflects a difference in the temporal shape of activity during exposure to the apparatus rather than a sustained decrease in locomotion.

## Discussion

This study provides a focused baseline behavioral characterization of male RORγt-Tg mice in the absence of an acute immune challenge. Two features were evident: lower body weight and an altered temporal profile of locomotor activity during initial exposure to the light–dark apparatus. In contrast, resident–intruder aggression and conventional light–dark box measures of anxiety-like behavior did not differ detectably between genotypes. The behavioral finding is therefore best interpreted as a difference in the dynamics of the initial response to a novel environment rather than as a broad change in anxiety-like behavior or overall locomotor capacity.

The body-weight phenotype was the strongest group difference in the present dataset. IL-17 signaling has previously been implicated in the regulation of adipogenesis, glucose homeostasis, and obesity-related phenotypes (Zúñiga et al., 2010), making the direction of the present effect biologically plausible in a line with a known Th17-biased immune phenotype. However, the present study does not establish that reduced body weight is mediated by IL-17A. Cytokine concentrations, Th17-cell abundance, food intake, and metabolic parameters were not measured in the tested animals. The body-weight result should therefore be viewed as a robust genotype-associated phenotype that may help guide future metabolic and immunological analyses of this line.

The locomotor time-course result is notable because it was dissociated from the conventional measures obtained in the same light–dark test. The light–dark box is commonly used as an approach–avoidance assay in which time spent in the illuminated compartment, transitions between compartments, and transition latency are interpreted as anxiety-related measures (Crawley and Goodwin, 1980; Bourin and Hascoët, 2003). None of these endpoints differed between genotypes. Instead, the difference emerged in how locomotor activity changed across time. Habituation to a novel environment is itself a measurable behavioral process that varies among mouse strains and can be dissociated from conventional anxiety-like measures (Leussis and Bolivar, 2006; Bolivar, 2009). Accordingly, the genotype × time interaction may reflect altered novelty responsiveness, early exploratory drive, or the kinetics of within-session habituation. Distinguishing among these possibilities will require dedicated paradigms designed to separate novelty detection, exploration, and habituation.

A mechanistic link to the RORγt–Th17–IL-17A axis remains an important hypothesis rather than a conclusion of the present experiment. The transgenic line is known to overexpress RORγt preferentially in T cells and to exhibit a Th17-biased phenotype with elevated circulating IL-17A (Yoh et al., 2012). Independent studies have shown that IL-17A can influence neuronal function, social behavior, anxiety-related behavior, and microglial responses (Choi et al., 2016; Alves de Lima et al., 2020; Reed et al., 2020; Sasaki et al., 2020). The present animals, however, were not phenotyped immunologically at the time of behavioral testing. Future studies that quantify circulating IL-17A and Th17-cell abundance in the same animals, or manipulate IL-17A/IL-17 receptor signaling, will be necessary to determine whether the altered locomotor trajectory depends on this pathway.

The absence of a detectable aggression phenotype should also be interpreted within the characteristics of the resident–intruder dataset. Overall attack rates were low, and no attacks occurred in either genotype during run 3. The resident–intruder paradigm is sensitive to housing, strain, experience, and procedural variables, and low baseline aggression can reduce power to detect genotype-dependent differences (Koolhaas et al., 2013). The current data therefore argue against a large alteration of offensive aggression under the conditions tested but do not exclude more subtle changes or effects that might emerge under a protocol optimized for higher baseline aggression.

Several limitations should be considered. The sample size was modest and restricted to males, and age varied across the three test runs. The minute-1 contrast was exploratory, was driven mainly by runs 2 and 3, and did not remain significant after correction across all ten bins; replication in an independent cohort is therefore warranted. In addition, the study did not measure Th17 cells or IL-17A in the behaviorally tested animals, preventing direct mechanistic attribution. Nevertheless, the significant genotype × time interaction, together with the robust body-weight difference and the absence of changes in conventional anxiety-like measures, defines a focused phenotype that can be tested in future mechanistic experiments. These findings establish a practical baseline for investigating how constitutive RORγt-dependent immune alterations may shape behavioral responses to novelty.

## Methods

### Animals

Male RORγt-Tg mice and WT littermates on a C57BL/6 background were used. The transgenic line corresponds to Tg(CD2-Rorc)#Staka (MGI:5805451), in which full-length murine RORγt cDNA is expressed under human CD2 regulatory elements, resulting in preferential RORγt overexpression in T cells (Yoh et al., 2012). The line was maintained as heterozygotes by backcrossing with C57BL/6 mice (Sasaki et al., 2021). Mice were tested in three runs and were 81–100 days old on the day of the light–dark box test (run 1: 3 WT/3 Tg, 100 days; run 2: 3 WT/3 Tg, 81 days; run 3: 6 WT/7 Tg, 93–99 days). Animals were maintained under specific pathogen-free conditions at the Laboratory Animal Resource Center, University of Tsukuba, under a 12:12-h light/dark cycle (lights on 07:00–19:00), with food and water available ad libitum. Animals received no treatment before behavioral testing. Genotypes were determined by PCR using an established colony-management protocol. All behavioral tests were conducted during the light phase. Mice were single-housed for 7 days (runs 1 and 2) or 9 days (run 3) before the first behavioral test and remained singly housed throughout testing.

All procedures were conducted in accordance with the Guidelines for the Care and Use of Laboratory Animals at the University of Tsukuba and were approved by the Animal Experiment Committee of the University of Tsukuba (approval nos. 24-151, 25-172, and 26-163). The use and breeding of genetically modified animals were approved by the University of Tsukuba Safety Committee for Recombinant DNA Experiments (approval no. 210262). Efforts were made to minimize animal suffering and the number of animals used.

### Resident–intruder test

Aggression was assessed on three consecutive days using the resident–intruder paradigm with modifications from a standardized protocol (Koolhaas et al., 2013). A bilaterally olfactory-bulbectomized ICR/Jcl male intruder (CLEA Japan, Inc., Tokyo, Japan) was introduced into the home cage of the singly housed resident, and the encounter was video-recorded for 15 min. Bulbectomy was used to reduce aggression initiated by the intruder so that scored attacks were attributable to the resident. Latency to the first resident attack bite and the total number of resident attack bites were scored offline from the video recordings. Residents that did not attack during a 15-min session were assigned a latency of 900 s. Attack latency was averaged across the three sessions, and attack bites were summed across the three sessions.

### Light–dark box test

On the day after the final resident–intruder session, each mouse was tested once in an O’Hara light–dark transition apparatus configured as described by Tsuda and Ogawa (2012). The apparatus consisted of an enclosed black dark compartment (0 lux) and an open-top white light compartment (350 lux), each measuring 40 × 20 × 25 cm, connected by a 5 × 2 cm doorway. At the start of the test, the mouse was placed in the dark compartment, and the doorway opened automatically after 5 s. Horizontal activity and compartment occupancy were recorded for 10 min on a Windows computer using Image J LD2 software (O’Hara & Co., Ltd., Tokyo, Japan). The software provided distance traveled in each compartment, cumulative time in the light compartment, latency to first light-compartment entry, and the number of transitions between compartments. Distance was also exported in 1-min bins; total distance for each bin was calculated as the sum of distance traveled in the dark and light compartments. After each test, mice were returned to their home cage and the apparatus was wiped clean.

### Body weight

Body weight was measured immediately after completion of the light–dark box test.

### Statistical analysis

Analyses were performed in Python 3.11.15 using SciPy 1.17.1 and Pingouin 0.6.1. All tests were two-tailed with α = 0.05, and data from the three runs were pooled (WT *n* = 12; Tg *n* = 13). Body weight, time in the light compartment, number of transitions, and latency to the first light-compartment entry were compared with Student’s two-sample *t*-tests. Attack latency and attack bite counts were compared with two-sided Mann–Whitney *U* tests using asymptotic *p*-values corrected for tied observations because latency was right-censored at the 900-s cut-off and bite counts were strongly zero-inflated. These endpoints are therefore summarized graphically by median and interquartile range. The proportion of residents that attacked at least once was compared with Fisher’s exact test.

Distance traveled in 1-min bins was analyzed with a mixed-model ANOVA with genotype as the between-subjects factor and time bin as the within-subjects factor. Mauchly’s test indicated a violation of sphericity (W = 0.023, *p* = 0.0015), so the Greenhouse–Geisser correction (ε = 0.537) was applied to the main effect of time and the genotype × time interaction, yielding corrected degrees of freedom of 4.83 and 111.10. The genotype main effect was between subjects and was reported without Greenhouse–Geisser correction. Bin-wise comparisons following the significant interaction were post hoc and exploratory; no analysis plan was preregistered. Unadjusted *p*-values are reported together with the Bonferroni-adjusted value across the ten bins for minute 1.

Sensitivity analyses used linear models including test run and age at testing as covariates for body weight and minute-1 distance. A separate model included body weight as a covariate for minute-1 distance. Unless otherwise specified, summary values are mean ± s.e.m. Figure generation used NumPy 2.4.4 and Matplotlib 3.10.9.

### AI-assisted manuscript preparation

ChatGPT (GPT-5.6 Sol, OpenAI) assisted with English-language editing, statistical-code review, figure and supplementary-data preparation, and manuscript formatting. No other AI tools were used. The authors critically reviewed all AI-assisted outputs against the original data and take full responsibility for the accuracy and integrity of the final manuscript.

## Data availability

The source data underlying all analyses and Figure 1, including per-animal behavioral and body-weight data, per-animal locomotor activity in 1-min bins, and the statistical-analysis and figure-generation code, are provided as Supplementary Data 1–3 accompanying this preprint and are also available from the corresponding authors upon reasonable request.

## Acknowledgements

We are grateful to Sae Sanaka and Kenyu Nakamura for valuable discussions and critical reading of this manuscript.

## Author contributions

Tetsuya Sasaki: Conceptualization, Data curation, Formal analysis, Funding acquisition, Investigation, Project administration, Supervision. Aki Takahashi: Data curation, Formal analysis, Methodology, Investigation. Yosuke Takei: Conceptualization, Project administration.

