## Supplementary Data 3 for "Male RORγt-transgenic mice exhibit lower body weight and an altered locomotor response to a novel environment": analysis_and_figure.py

```
#!/usr/bin/env python3
"""
Reproduce the statistical analyses and data-derived panels for the RORγt-Tg
behavioral microPublication.

Inputs
------
Extended_Data_1_Per_animal_behavioral_and_body_weight_data.csv
Extended_Data_2_Per_animal_locomotor_activity_1min_bins.csv

Outputs
-------
analysis_results.txt
Figure1_reproduced.png

The script uses only NumPy, SciPy, and Matplotlib in addition to the Python
standard library.
"""

from __future__ import annotations

import argparse
import csv
import math
from pathlib import Path
from typing import Dict, Iterable, List, Sequence, Tuple

import matplotlib.pyplot as plt
from matplotlib.gridspec import GridSpec
from matplotlib.patches import FancyArrowPatch, Rectangle
import numpy as np
from scipy import stats


def read_summary(path: Path) -> List[dict]:
    rows: List[dict] = []
    with path.open("r", encoding="utf-8-sig", newline="") as handle:
        for raw in csv.DictReader(handle):
            rows.append(
                {
                    "mouse_id": int(raw["mouse_id"]),
                    "sex": raw["sex"],
                    "genotype": raw["genotype"],
                    "test_run": int(raw["test_run"]),
                    "age_days": int(raw["age_days"]),
                    "age_weeks": float(raw["age_weeks"]),
                    "single_housing_days": int(raw["single_housing_days"]),
                    "body_weight_g": float(raw["body_weight_g"]),
                    "attack_latency_mean_s": float(raw["attack_latency_mean_s"]),
                    "attack_bites_total": int(raw["attack_bites_total"]),
                    "attacked_at_least_once": raw["attacked_at_least_once"].strip().lower()
                    in {"true", "1", "yes"},
                    "time_in_light_s": float(raw["time_in_light_s"]),
                    "transitions_n": int(raw["transitions_n"]),
                    "latency_to_first_light_transition_s": float(
                        raw["latency_to_first_light_transition_s"]
                    ),
                }
            )
    return rows


def read_locomotor(path: Path) -> List[dict]:
    rows: List[dict] = []
    with path.open("r", encoding="utf-8-sig", newline="") as handle:
        for raw in csv.DictReader(handle):
            rows.append(
                {
                    "mouse_id": int(raw["mouse_id"]),
                    "sex": raw["sex"],
                    "genotype": raw["genotype"],
                    "test_run": int(raw["test_run"]),
                    "age_days": int(raw["age_days"]),
                    "minute_bin": int(raw["minute_bin"]),
                    "dark_distance_cm": float(raw["dark_distance_cm"]),
                    "light_distance_cm": float(raw["light_distance_cm"]),
                    "total_distance_cm": float(raw["total_distance_cm"]),
                }
            )
    return rows


def group_values(rows: Sequence[dict], key: str, genotype: str) -> np.ndarray:
    return np.asarray([float(r[key]) for r in rows if r["genotype"] == genotype])


def mean_sem(values: Sequence[float]) -> Tuple[float, float]:
    x = np.asarray(values, dtype=float)
    return float(x.mean()), float(x.std(ddof=1) / math.sqrt(x.size))


def ols_table(y: Sequence[float], x: np.ndarray) -> dict:
    y_arr = np.asarray(y, dtype=float)
    x_arr = np.asarray(x, dtype=float)
    beta = np.linalg.lstsq(x_arr, y_arr, rcond=None)[0]
    residuals = y_arr - x_arr @ beta
    rank = np.linalg.matrix_rank(x_arr)
    df = y_arr.size - rank
    mse = float(np.sum(residuals**2) / df)
    covariance = mse * np.linalg.pinv(x_arr.T @ x_arr)
    se = np.sqrt(np.diag(covariance))
    t_values = beta / se
    p_values = 2 * stats.t.sf(np.abs(t_values), df)
    return {
        "beta": beta,
        "se": se,
        "t": t_values,
        "p": p_values,
        "df": int(df),
    }


def wide_locomotor(locomotor: Sequence[dict]) -> Tuple[List[int], np.ndarray, np.ndarray]:
    mouse_ids = sorted({r["mouse_id"] for r in locomotor})
    matrix = np.zeros((len(mouse_ids), 10), dtype=float)
    genotype = []
    for i, mouse_id in enumerate(mouse_ids):
        mouse_rows = [r for r in locomotor if r["mouse_id"] == mouse_id]
        by_minute = {r["minute_bin"]: r for r in mouse_rows}
        if set(by_minute) != set(range(1, 11)):
            raise ValueError(f"Mouse {mouse_id} does not have exactly ten minute bins.")
        matrix[i, :] = [by_minute[m]["total_distance_cm"] for m in range(1, 11)]
        genotype.append(mouse_rows[0]["genotype"])
    return mouse_ids, matrix, np.asarray(genotype)


def greenhouse_geisser_epsilon(matrix: np.ndarray) -> float:
    covariance = np.cov(matrix, rowvar=False, ddof=1)
    k = covariance.shape[0]
    mean_variance = np.diag(covariance).mean()
    grand_covariance_mean = covariance.mean()
    ss_matrix = np.sum(covariance**2)
    ss_rows = np.sum(covariance.mean(axis=1) ** 2)
    numerator = (k * (mean_variance - grand_covariance_mean)) ** 2
    denominator = (k - 1) * (
        ss_matrix
        - 2 * k * ss_rows
        + (k**2) * (grand_covariance_mean**2)
    )
    return float(min(numerator / denominator, 1.0))


def mauchly_sphericity(matrix: np.ndarray) -> dict:
    n, k = matrix.shape
    d = k - 1
    covariance = np.cov(matrix, rowvar=False, ddof=1)
    centered = (
        covariance
        - covariance.mean(axis=0)[:, None]
        - covariance.mean(axis=1)[None, :]
        + covariance.mean()
    )
    eigenvalues = np.linalg.eigvalsh(centered)[1:]
    tolerance = np.finfo(float).eps * eigenvalues.max() * d
    eigenvalues = eigenvalues[eigenvalues > tolerance]
    w_value = float(np.prod(eigenvalues) / (eigenvalues.sum() / d) ** d)
    log_w = math.log(w_value)
    factor = 1 - (2 * d**2 + d + 2) / (6 * d * (n - 1))
    correction = (
        (d + 2)
        * (d - 1)
        * (d - 2)
        * (2 * d**3 + 6 * d**2 + 3 * k + 2)
        / (288 * ((n - 1) * d * factor) ** 2)
    )
    chi_square = -(n - 1) * factor * log_w
    df = int((d * (d + 1)) / 2 - 1)
    p1 = stats.chi2.sf(chi_square, df)
    p2 = stats.chi2.sf(chi_square, df + 4)
    p_value = float(p1 + correction * (p2 - p1))
    return {"W": w_value, "chi2": float(chi_square), "df": df, "p": p_value}


def mixed_anova(matrix: np.ndarray, genotype: np.ndarray) -> dict:
    n_subjects, n_times = matrix.shape
    groups = ["WT", "Tg"]
    grand_mean = matrix.mean()
    subject_means = matrix.mean(axis=1)
    time_means = matrix.mean(axis=0)

    ss_between = n_times * np.sum((subject_means - grand_mean) ** 2)
    group_means = {g: matrix[genotype == g].mean() for g in groups}
    ss_group = n_times * sum(
        np.sum(genotype == g) * (group_means[g] - grand_mean) ** 2 for g in groups
    )
    ss_subject_group = ss_between - ss_group

    ss_within = np.sum((matrix - subject_means[:, None]) ** 2)
    ss_time = n_subjects * np.sum((time_means - grand_mean) ** 2)
    ss_interaction = 0.0
    for group in groups:
        n_group = np.sum(genotype == group)
        group_time_means = matrix[genotype == group].mean(axis=0)
        ss_interaction += n_group * np.sum(
            (
                group_time_means
                - group_means[group]
                - time_means
                + grand_mean
            )
            ** 2
        )
    ss_error = ss_within - ss_time - ss_interaction

    df_group = 1
    df_subject_group = n_subjects - len(groups)
    df_time = n_times - 1
    df_interaction = df_time
    df_error = df_subject_group * df_time

    f_group = (ss_group / df_group) / (ss_subject_group / df_subject_group)
    f_time = (ss_time / df_time) / (ss_error / df_error)
    f_interaction = (ss_interaction / df_interaction) / (ss_error / df_error)

    epsilon = greenhouse_geisser_epsilon(matrix)
    corrected_df1 = df_time * epsilon
    corrected_df2 = df_error * epsilon

    return {
        "F_group": float(f_group),
        "df_group_1": df_group,
        "df_group_2": df_subject_group,
        "p_group": float(stats.f.sf(f_group, df_group, df_subject_group)),
        "F_time": float(f_time),
        "F_interaction": float(f_interaction),
        "df_time_uncorrected_1": df_time,
        "df_time_uncorrected_2": df_error,
        "epsilon_GG": epsilon,
        "df_GG_1": float(corrected_df1),
        "df_GG_2": float(corrected_df2),
        "p_time_GG": float(stats.f.sf(f_time, corrected_df1, corrected_df2)),
        "p_interaction_GG": float(
            stats.f.sf(f_interaction, corrected_df1, corrected_df2)
        ),
    }


def analysis(summary: Sequence[dict], locomotor: Sequence[dict]) -> dict:
    output: Dict[str, object] = {}

    for key in [
        "body_weight_g",
        "time_in_light_s",
        "transitions_n",
        "latency_to_first_light_transition_s",
    ]:
        wt = group_values(summary, key, "WT")
        tg = group_values(summary, key, "Tg")
        test = stats.ttest_ind(wt, tg, equal_var=True)
        output[key] = {
            "t": float(test.statistic),
            "df": int(wt.size + tg.size - 2),
            "p": float(test.pvalue),
        }

    for key in ["attack_latency_mean_s", "attack_bites_total"]:
        wt = group_values(summary, key, "WT")
        tg = group_values(summary, key, "Tg")
        test = stats.mannwhitneyu(
            wt, tg, alternative="two-sided", method="asymptotic"
        )
        output[key] = {"U": float(test.statistic), "p": float(test.pvalue)}

    wt_attacked = sum(
        bool(r["attacked_at_least_once"])
        for r in summary
        if r["genotype"] == "WT"
    )
    tg_attacked = sum(
        bool(r["attacked_at_least_once"])
        for r in summary
        if r["genotype"] == "Tg"
    )
    contingency = np.asarray(
        [
            [wt_attacked, 12 - wt_attacked],
            [tg_attacked, 13 - tg_attacked],
        ]
    )
    fisher = stats.fisher_exact(contingency, alternative="two-sided")
    output["attack_incidence"] = {
        "WT_attacked": int(wt_attacked),
        "WT_total": 12,
        "Tg_attacked": int(tg_attacked),
        "Tg_total": 13,
        "odds_ratio": float(fisher.statistic),
        "p": float(fisher.pvalue),
    }

    mouse_ids, matrix, genotype = wide_locomotor(locomotor)
    output["mauchly"] = mauchly_sphericity(matrix)
    output["mixed_anova"] = mixed_anova(matrix, genotype)

    minute_tests = []
    for minute in range(1, 11):
        wt = np.asarray(
            [
                r["total_distance_cm"]
                for r in locomotor
                if r["genotype"] == "WT" and r["minute_bin"] == minute
            ]
        )
        tg = np.asarray(
            [
                r["total_distance_cm"]
                for r in locomotor
                if r["genotype"] == "Tg" and r["minute_bin"] == minute
            ]
        )
        test = stats.ttest_ind(wt, tg, equal_var=True)
        minute_tests.append(
            {
                "minute": minute,
                "t": float(test.statistic),
                "df": int(wt.size + tg.size - 2),
                "p_unadjusted": float(test.pvalue),
                "p_bonferroni": float(min(1.0, test.pvalue * 10)),
            }
        )
    output["minute_tests"] = minute_tests

    summary_by_id = {r["mouse_id"]: r for r in summary}
    ordered_ids = sorted(summary_by_id)
    genotype_num = np.asarray(
        [1 if summary_by_id[m]["genotype"] == "Tg" else 0 for m in ordered_ids],
        dtype=float,
    )
    run = np.asarray([summary_by_id[m]["test_run"] for m in ordered_ids])
    age = np.asarray([summary_by_id[m]["age_days"] for m in ordered_ids], dtype=float)
    run_age_design = np.column_stack(
        [
            np.ones(len(ordered_ids)),
            genotype_num,
            (run == 2).astype(float),
            (run == 3).astype(float),
            age - age.mean(),
        ]
    )
    body_weight = np.asarray(
        [summary_by_id[m]["body_weight_g"] for m in ordered_ids], dtype=float
    )
    bin1 = {
        r["mouse_id"]: r["total_distance_cm"]
        for r in locomotor
        if r["minute_bin"] == 1
    }
    bin1_values = np.asarray([bin1[m] for m in ordered_ids], dtype=float)

    body_model = ols_table(body_weight, run_age_design)
    bin1_model = ols_table(bin1_values, run_age_design)
    weight_design = np.column_stack([np.ones(len(ordered_ids)), genotype_num, body_weight])
    weight_model = ols_table(bin1_values, weight_design)
    output["sensitivity"] = {
        "body_weight_genotype_p_run_age_adjusted": float(body_model["p"][1]),
        "bin1_genotype_p_run_age_adjusted": float(bin1_model["p"][1]),
        "bin1_genotype_p_weight_adjusted": float(weight_model["p"][1]),
        "bin1_body_weight_p": float(weight_model["p"][2]),
    }
    return output


RUN_MARKERS = {1: "o", 2: "^", 3: "s"}
GROUP_COLORS = {"WT": "#0072B2", "Tg": "#D50000"}
GROUP_FACE = {"WT": "#CFE8F3", "Tg": "#F4CCCC"}


def points_for_group(ax, rows: Sequence[dict], key: str, genotype: str, x: float) -> None:
    group_rows = [r for r in rows if r["genotype"] == genotype]
    offsets = np.linspace(-0.12, 0.12, len(group_rows))
    for offset, row in zip(offsets, group_rows):
        ax.scatter(
            x + offset,
            row[key],
            marker=RUN_MARKERS[row["test_run"]],
            facecolors="white",
            edgecolors=GROUP_COLORS[genotype],
            linewidths=1.2,
            s=28,
            zorder=4,
        )


def bar_panel(ax, rows: Sequence[dict], key: str, ylabel: str, title: str, stat_text: str) -> None:
    groups = ["WT", "Tg"]
    for x, group in enumerate(groups):
        values = group_values(rows, key, group)
        mean, sem = mean_sem(values)
        ax.bar(
            x,
            mean,
            yerr=sem,
            width=0.58,
            color=GROUP_FACE[group],
            edgecolor=GROUP_COLORS[group],
            linewidth=1.5,
            capsize=4,
            zorder=2,
        )
        points_for_group(ax, rows, key, group, x)
    ax.set_xticks([0, 1], ["WT", "Tg"])
    ax.set_ylabel(ylabel)
    ax.set_title(title, fontsize=10)
    ax.spines[["top", "right"]].set_visible(False)
    ax.text(0.5, -0.28, stat_text, transform=ax.transAxes, ha="center", fontsize=8)


def box_panel(ax, rows: Sequence[dict], key: str, ylabel: str, title: str, stat_text: str) -> None:
    groups = ["WT", "Tg"]
    values = [group_values(rows, key, g) for g in groups]
    bp = ax.boxplot(
        values,
        positions=[0, 1],
        widths=0.55,
        patch_artist=True,
        showfliers=False,
        medianprops={"color": "black", "linewidth": 1.5},
        whiskerprops={"linewidth": 1.2},
        capprops={"linewidth": 1.2},
    )
    for patch, group in zip(bp["boxes"], groups):
        patch.set_facecolor(GROUP_FACE[group])
        patch.set_edgecolor(GROUP_COLORS[group])
        patch.set_linewidth(1.5)
    for x, group in enumerate(groups):
        points_for_group(ax, rows, key, group, x)
    ax.set_xticks([0, 1], ["WT", "Tg"])
    ax.set_ylabel(ylabel)
    ax.set_title(title, fontsize=10)
    ax.spines[["top", "right"]].set_visible(False)
    ax.text(0.5, -0.28, stat_text, transform=ax.transAxes, ha="center", fontsize=8)


def panel_label(ax, label: str) -> None:
    ax.text(-0.25, 1.12, label, transform=ax.transAxes, fontsize=15, fontweight="bold")


def plot_figure(summary: Sequence[dict], locomotor: Sequence[dict], results: dict, output: Path) -> None:
    fig = plt.figure(figsize=(12, 9), dpi=300)
    grid = GridSpec(3, 4, figure=fig, height_ratios=[0.75, 1.0, 1.0], hspace=0.85, wspace=0.55)

    # Panel A: schematic timeline
    ax_a = fig.add_subplot(grid[0, :])
    ax_a.set_xlim(0, 1)
    ax_a.set_ylim(0, 1)
    ax_a.axis("off")
    panel_label(ax_a, "A")
    blocks = [
        (0.08, 0.58, 0.18, 0.24, "Single housing", "7–9 days"),
        (0.41, 0.58, 0.22, 0.24, "Resident–intruder test", "15 min/day × 3 days"),
        (0.77, 0.58, 0.18, 0.24, "Light–dark box test", "10 min, then body weight"),
    ]
    for x, y, w, h, heading, sub in blocks:
        ax_a.add_patch(Rectangle((x, y), w, h, facecolor="#F3F4F6", edgecolor="#555555", linewidth=1.2))
        ax_a.text(x + w / 2, y + h + 0.08, heading, ha="center", fontsize=10, fontweight="bold")
        ax_a.text(x + w / 2, y - 0.10, sub, ha="center", fontsize=8)
    for start, end in [((0.27, 0.70), (0.40, 0.70)), ((0.64, 0.70), (0.76, 0.70))]:
        ax_a.add_patch(FancyArrowPatch(start, end, arrowstyle="-|>", mutation_scale=12, color="#777777", linewidth=1.2))
    ax_a.plot([0.06, 0.96], [0.22, 0.22], color="#AAAAAA", linewidth=1)
    ax_a.text(0.17, 0.08, "Day −9 to −7", ha="center", fontsize=8)
    ax_a.text(0.52, 0.08, "Day 0–2", ha="center", fontsize=8)
    ax_a.text(0.86, 0.08, "Day 3", ha="center", fontsize=8)

    ax_b = fig.add_subplot(grid[1, 0])
    bar_panel(
        ax_b, summary, "body_weight_g", "Body weight (g)", "Body weight",
        f"t(23) = {results['body_weight_g']['t']:.2f}, p = {results['body_weight_g']['p']:.3f}"
    )
    panel_label(ax_b, "B")

    ax_c = fig.add_subplot(grid[1, 1])
    box_panel(
        ax_c, summary, "attack_latency_mean_s", "Attack latency (s)", "Resident–intruder: latency",
        f"U = {results['attack_latency_mean_s']['U']:.0f}, p = {results['attack_latency_mean_s']['p']:.3f}"
    )
    ax_c.axhline(900, color="#777777", linestyle="--", linewidth=0.8)
    panel_label(ax_c, "C")

    ax_d = fig.add_subplot(grid[1, 2])
    box_panel(
        ax_d, summary, "attack_bites_total", "Attack bites (n)", "Resident–intruder: bites",
        f"U = {results['attack_bites_total']['U']:.0f}, p = {results['attack_bites_total']['p']:.3f}"
    )
    panel_label(ax_d, "D")

    ax_e = fig.add_subplot(grid[1, 3])
    bar_panel(
        ax_e, summary, "time_in_light_s", "Time in light (s)", "Light–dark: time in light",
        f"t(23) = {results['time_in_light_s']['t']:.2f}, p = {results['time_in_light_s']['p']:.3f}"
    )
    panel_label(ax_e, "E")

    ax_f = fig.add_subplot(grid[2, 0])
    bar_panel(
        ax_f, summary, "transitions_n", "Transitions (n)", "Light–dark: transitions",
        f"t(23) = {results['transitions_n']['t']:.2f}, p = {results['transitions_n']['p']:.3f}"
    )
    panel_label(ax_f, "F")

    ax_g = fig.add_subplot(grid[2, 1])
    bar_panel(
        ax_g, summary, "latency_to_first_light_transition_s",
        "Latency to first transition (s)", "Light–dark: transition latency",
        f"t(23) = {results['latency_to_first_light_transition_s']['t']:.2f}, p = {results['latency_to_first_light_transition_s']['p']:.3f}"
    )
    panel_label(ax_g, "G")

    ax_h = fig.add_subplot(grid[2, 2:])
    for genotype, marker in [("WT", "o"), ("Tg", "s")]:
        means = []
        sems = []
        for minute in range(1, 11):
            values = [
                r["total_distance_cm"]
                for r in locomotor
                if r["genotype"] == genotype and r["minute_bin"] == minute
            ]
            mean, sem = mean_sem(values)
            means.append(mean)
            sems.append(sem)
        ax_h.errorbar(
            range(1, 11),
            means,
            yerr=sems,
            marker=marker,
            linewidth=1.5,
            capsize=3,
            color=GROUP_COLORS[genotype],
            markerfacecolor="white",
            markeredgewidth=1.5,
            label=f"{genotype} (n = {12 if genotype == 'WT' else 13})",
        )
    ax_h.set_xlabel("Time bin (min)")
    ax_h.set_ylabel("Distance traveled\nper 1-min bin (cm)")
    ax_h.set_xticks(range(1, 11))
    ax_h.set_title("Light–dark box: locomotor time course", fontsize=10)
    ax_h.legend(frameon=False, fontsize=8)
    ax_h.spines[["top", "right"]].set_visible(False)
    mixed = results["mixed_anova"]
    ax_h.text(
        0.02,
        0.03,
        (
            f"Genotype × time: F_GG({mixed['df_GG_1']:.2f}, {mixed['df_GG_2']:.2f}) "
            f"= {mixed['F_interaction']:.2f}, p = {mixed['p_interaction_GG']:.3f}\n"
            f"Time: F_GG({mixed['df_GG_1']:.2f}, {mixed['df_GG_2']:.2f}) "
            f"= {mixed['F_time']:.2f}, p < 0.001; "
            f"genotype: F(1, 23) = {mixed['F_group']:.2f}, p = {mixed['p_group']:.3f}"
        ),
        transform=ax_h.transAxes,
        fontsize=7,
        va="bottom",
    )
    panel_label(ax_h, "H")

    fig.subplots_adjust(left=0.07, right=0.98, top=0.96, bottom=0.08)
    fig.savefig(output, dpi=300)
    plt.close(fig)


def write_results(results: dict, path: Path) -> None:
    lines = []
    lines.append("PRIMARY BETWEEN-GENOTYPE COMPARISONS")
    lines.append(
        f"Body weight: t(23) = {results['body_weight_g']['t']:.6f}, "
        f"p = {results['body_weight_g']['p']:.9f}"
    )
    lines.append(
        f"Attack latency: U = {results['attack_latency_mean_s']['U']:.0f}, "
        f"p = {results['attack_latency_mean_s']['p']:.9f}"
    )
    lines.append(
        f"Attack bites: U = {results['attack_bites_total']['U']:.0f}, "
        f"p = {results['attack_bites_total']['p']:.9f}"
    )
    incidence = results["attack_incidence"]
    lines.append(
        f"Attack incidence: {incidence['WT_attacked']}/{incidence['WT_total']} WT vs "
        f"{incidence['Tg_attacked']}/{incidence['Tg_total']} Tg; "
        f"Fisher p = {incidence['p']:.9f}"
    )
    lines.append(
        f"Time in light: t(23) = {results['time_in_light_s']['t']:.6f}, "
        f"p = {results['time_in_light_s']['p']:.9f}"
    )
    lines.append(
        f"Transitions: t(23) = {results['transitions_n']['t']:.6f}, "
        f"p = {results['transitions_n']['p']:.9f}"
    )
    lines.append(
        f"Transition latency: t(23) = "
        f"{results['latency_to_first_light_transition_s']['t']:.6f}, "
        f"p = {results['latency_to_first_light_transition_s']['p']:.9f}"
    )
    lines.append("")
    lines.append("SPHERICITY AND MIXED ANOVA")
    mauchly = results["mauchly"]
    lines.append(
        f"Mauchly: W = {mauchly['W']:.9f}, chi2({mauchly['df']}) = "
        f"{mauchly['chi2']:.6f}, p = {mauchly['p']:.9f}"
    )
    mixed = results["mixed_anova"]
    lines.append(f"Greenhouse–Geisser epsilon = {mixed['epsilon_GG']:.9f}")
    lines.append(
        f"Genotype × time: F_GG({mixed['df_GG_1']:.6f}, {mixed['df_GG_2']:.6f}) "
        f"= {mixed['F_interaction']:.6f}, p = {mixed['p_interaction_GG']:.9f}"
    )
    lines.append(
        f"Time: F_GG({mixed['df_GG_1']:.6f}, {mixed['df_GG_2']:.6f}) "
        f"= {mixed['F_time']:.6f}, p = {mixed['p_time_GG']:.12g}"
    )
    lines.append(
        f"Genotype: F(1, 23) = {mixed['F_group']:.6f}, p = {mixed['p_group']:.9f}"
    )
    lines.append("")
    lines.append("BIN-WISE POST HOC TESTS")
    for row in results["minute_tests"]:
        lines.append(
            f"Minute {row['minute']}: t({row['df']}) = {row['t']:.6f}, "
            f"unadjusted p = {row['p_unadjusted']:.9f}, "
            f"Bonferroni p = {row['p_bonferroni']:.9f}"
        )
    lines.append("")
    lines.append("SENSITIVITY ANALYSES")
    sens = results["sensitivity"]
    lines.append(
        f"Body-weight genotype term adjusted for run and age: "
        f"p = {sens['body_weight_genotype_p_run_age_adjusted']:.9f}"
    )
    lines.append(
        f"Minute-1 genotype term adjusted for run and age: "
        f"p = {sens['bin1_genotype_p_run_age_adjusted']:.9f}"
    )
    lines.append(
        f"Minute-1 genotype term adjusted for body weight: "
        f"p = {sens['bin1_genotype_p_weight_adjusted']:.9f}"
    )
    lines.append(
        f"Body-weight term in minute-1 model: p = {sens['bin1_body_weight_p']:.9f}"
    )
    path.write_text("\n".join(lines) + "\n", encoding="utf-8")


def main() -> None:
    parser = argparse.ArgumentParser()
    parser.add_argument(
        "--summary-csv",
        type=Path,
        default=Path("Extended_Data_1_Per_animal_behavioral_and_body_weight_data.csv"),
    )
    parser.add_argument(
        "--locomotor-csv",
        type=Path,
        default=Path("Extended_Data_2_Per_animal_locomotor_activity_1min_bins.csv"),
    )
    parser.add_argument("--output-dir", type=Path, default=Path("analysis_output"))
    args = parser.parse_args()

    args.output_dir.mkdir(parents=True, exist_ok=True)
    summary = read_summary(args.summary_csv)
    locomotor = read_locomotor(args.locomotor_csv)
    results = analysis(summary, locomotor)
    write_results(results, args.output_dir / "analysis_results.txt")
    plot_figure(summary, locomotor, results, args.output_dir / "Figure1_reproduced.png")
    print(f"Analysis complete: {args.output_dir.resolve()}")


if __name__ == "__main__":
    main()
```
