## Supplementary Data 3 for "Male RORγt-transgenic mice exhibit lower body weight and an altered locomotor response to a novel environment": README.md

# Extended Data 3. Statistical-analysis and figure-generation code

**Resource type:** Software

**Brief description:** Python source code that reproduces the statistical analyses
reported in the manuscript and generates a 4:3 reproduction of the data-derived
Figure 1 panels from Extended Data 1 and Extended Data 2.

## Files

- `analysis_and_figure.py` — analysis and plotting script.
- `requirements.txt` — Python package requirements.
- `EXPECTED_RESULTS.txt` — reference output generated from the submitted data.
- `LICENSE` — MIT License for the analysis code.

## Input files

Place the following two CSV files in the same working directory as the script:

1. `Extended_Data_1_Per_animal_behavioral_and_body_weight_data.csv`
2. `Extended_Data_2_Per_animal_locomotor_activity_1min_bins.csv`

## Running the analysis

```bash
python -m venv .venv
# Linux/macOS
source .venv/bin/activate
# Windows PowerShell
# .venv\Scripts\Activate.ps1

pip install -r requirements.txt

python analysis_and_figure.py \
  --summary-csv Extended_Data_1_Per_animal_behavioral_and_body_weight_data.csv \
  --locomotor-csv Extended_Data_2_Per_animal_locomotor_activity_1min_bins.csv \
  --output-dir analysis_output
```

The script creates:

- `analysis_output/analysis_results.txt`
- `analysis_output/Figure1_reproduced.png`

## Statistical implementation

The script performs Student's two-sample t-tests, asymptotic two-sided
Mann–Whitney U tests with tie correction, Fisher's exact test, a two-way mixed
ANOVA, Mauchly's test, Greenhouse–Geisser correction, bin-wise t-tests with
Bonferroni correction, and the sensitivity linear models described in the
manuscript.

## License

The code is released under the MIT License. The data files are released under
CC BY 4.0.
